# Pericyte Loss Reprogrammes Capillary Endothelium and Drives White Matter Injury in Small Vessel Disease

**DOI:** 10.64898/2026.05.11.724300

**Authors:** D. Stefancova, A. Chagnot, M. Sewell, N. Fialova, C. McQuaid, O.J. Uweru, S. Becker, M. Romero-Bernal, W. Mungall, V. Walczak-Gillies, T.D. Farr, R. Lennen, M.A. Jansen, O. Dando, J. Cholewa-Waclaw, A. Montagne

**Author notes:** Contributed equally (D. Stefancova, A. Chagnot, M. Sewell).

## Abstract

Pericytes are critical regulators of cerebrovascular homeostasis, yet their contribution to small vessel disease (SVD) and white matter injury remains incompletely understood. Here, we use an inducible *Atp13a5*-driven genetic strategy to selectively label and ablate brain pericytes, enabling integrated morphological, functional, and transcriptomic analyses across the neurogliovascular unit. Single-cell RNA sequencing and tissue-level mapping identified distinct pericyte subtypes distributed along the vascular tree and revealed subtype-specific vulnerability following depletion. Moderate pericyte loss induced transient cerebrovascular dysfunction characterised by reduced cerebral blood flow and increased blood-brain barrier (BBB) permeability, accompanied by delayed white matter abnormalities, including altered diffusion MRI metrics, oligodendrocyte progenitor cell responses, and myelin loss. At the molecular level, pericyte depletion reprogrammed capillary endothelial cells toward an activated venular-like state characterised by VCAM-1 induction, reduced expression of the BBB-associated transporter MFSD2A, and activation of type I interferon signalling pathways. Cross-species analyses revealed enrichment of human white matter hyperintensity-associated SVD gene signatures across endothelial subtypes following pericyte depletion. Together, these findings identify pericyte dysfunction as a driver of endothelial inflammatory remodelling and white matter injury and establish mechanistic links between microvascular pericyte loss and human SVD.

## Introduction

Cerebral small vessel disease (SVD) is one of the leading causes of vascular cognitive impairment and dementia and contributes substantially to stroke, white matter degeneration, and age-related cognitive decline^1^. Hallmark neuroimaging features of SVD include white matter hyperintensities (WMH), enlarged perivascular spaces (PVS), lacunes, cerebral microbleeds, and brain atrophy, all of which are thought to arise, at least in part, from chronic dysfunction of the cerebral microvasculature. Increasing evidence further indicates that vascular dysfunction is not restricted to classical vascular dementias but also contributes to the pathophysiology of Alzheimer’s disease (AD) and other neurodegenerative disorders^2,3^. However, despite growing recognition of the importance of the neurogliovascular unit (NGVU) in brain ageing and dementia, the cellular and molecular mechanisms linking microvascular dysfunction to white matter injury and neurodegeneration remain incompletely understood.

Pericytes are mural cells embedded within the vascular basement membrane that closely interact with endothelial cells throughout the capillary network. Within the central nervous system (CNS), pericytes are increasingly recognised as critical regulators of cerebral blood flow (CBF), blood-brain barrier (BBB) integrity, angiogenesis, immune signalling, and vascular stability^4–6^. Pericyte dysfunction and loss have been reported in ageing, AD, and SVD, where they are associated with BBB leakage, hypoperfusion, and white matter damage^7–11^. A recent single-nuclei RNA-sequencing systematic review from our group further suggests that pericytes and endothelial cells are among the vascular cell populations most transcriptionally altered in AD and SVD, including enrichment of several disease-associated Genome-Wide Association Study (GWAS) genes within these vascular compartments^12^. In parallel, we studied human postmortem samples and demonstrated that pericyte loss occurs progressively across SVD severity and is associated with capillary endothelial remodelling characterised by inflammatory activation and a partial venular-like endothelial identity, including increased vascular cell adhesion molecule-1 (VCAM-1) expression^11^. Whether such endothelial changes can be recapitulated experimentally following selective pericyte dysfunction, and which molecular pathways drive these transitions, remain important unresolved questions.

One major limitation in the field has been the lack of highly selective tools to genetically manipulate brain pericytes. Commonly used mural cell markers, including *Pdgfrb*/PDGFRβ, *Cspg4*/NG2, and *Anpep*/CD13, are not restricted to capillary pericytes and can also label vascular smooth muscle cells (VSMCs), fibroblast-like perivascular cells, or peripheral populations^6,13^. Single-cell transcriptomic studies have further highlighted the remarkable heterogeneity of mural cell and endothelial cell populations along the arteriovenous axis, suggesting that pericytes exist along a transcriptional and anatomical continuum rather than as discrete cell classes^13,14^. Morphologically, brain pericytes have been subdivided into ensheathing, mesh, and thin-strand populations distributed across distinct vascular territories^15^. However, the physiological functions and disease susceptibilities of these morphotypes remain poorly understood.

Recent work identified *Atp13a5* as one of the most CNS-enriched pericyte markers currently available, with highly restricted expression within the brain vasculature^16^. This model therefore offers a unique opportunity to selectively investigate the role of brain pericytes while minimising off-target recombination of peripheral mural cell populations. In parallel, advances in high-resolution imaging, machine learning (ML)-based image analysis, longitudinal MRI, and single-cell RNA sequencing (scRNA-seq) now provide unprecedented opportunities to study pericyte heterogeneity and vascular dysfunction across scales, from microvascular morphology to transcriptomic remodelling.

In the present study, we combined an *Atp13a5*-driven inducible depletion strategy with advanced microscopy, ML-based morphometric analyses, longitudinal MRI, and scRNA-seq to investigate the consequences of moderate brain pericyte loss on cerebrovascular function, white matter integrity, and endothelial cell states. We demonstrate that selective pericyte depletion induces transient reductions in CBF, BBB dysfunction, white matter abnormalities, and marked endothelial inflammatory remodelling characterised by interferon-associated signalling, VCAM-1 induction, and loss of BBB-associated programmes. Finally, we show that endothelial transcriptional changes following pericyte depletion overlap with gene signatures associated with human WMH-related SVD, supporting the concept that moderate pericyte dysfunction may contribute to vascular states relevant to human SVD and white matter injury.

## Materials and Methods

### Animals

Mice were housed in ventilated plastic cages in groups of four to five with a 12-hour light/dark cycle. Animals had *ad libitum* access to water and a standard chow diet. All animals were bred and maintained in the animal facility at the BioQuarter, the University of Edinburgh. Animal care and experimental procedures were carried out in accordance with ARRIVE and UK Home Office regulations, under project licence PP7610194 (Cerebrovascular and inflamm-ageing link to neurodegeneration and dementia). Environmental enrichment in the form of nesting material and shelter was provided to promote natural behaviour and improve animal welfare.

For all experiments, sample sizes were determined through power calculations conducted prior to the start of the study. Male and female mice aged 5-7-month-old were used, except for single-cell RNA-sequencing experiments, in which mice were 9-12-month-old. All experiments were performed blinded, and no adverse events were observed.

### Generation and colony maintenance of Atp13a5-2A-CreERT2-IRES-tdTomato knock-in animal model

The *Atp13a5*^*tdT*^ transgenic mouse model was generated as previously described by Guo et al., 2024^16^. Sperm was collected from homozygous transgenic male mice at the University of Southern California in accordance with the guidelines of the United States National Institute of Health. The collected sperm samples were cryopreserved and transported to Edinburgh, United Kingdom.

In the animal facility at the University of Edinburgh, oocytes from healthy C57BL/6J female mice were harvested following a standard superovulation protocol. The collected oocytes were fertilised using thawed sperm, and resulting embryos were transferred into pseudopregnant female mice. Offspring were genotyped by Transnetyx (Cordova,TN) to confirm the presence of the knock-in allele.

Heterozygous mice were interbred to establish a homozygous colony. These homozygous *Atp13a5*^*tdT*^ transgenic mice were subsequently crossed with *iDTR* transgenic mice to generate the *Atp13a5;iDTR* mouse line, which was used to explore short- and long-term consequences of brain pericyte ablation.

### Genotyping of Atp13a5-2A-CreERT2-IRES-tdTomato knock-in model

The mouse genotype was determined from ear biopsies collected by animal technicians at the University of Edinburgh and shipped to Transnetyx (Cordova, TN) for DNA extraction and PCR-based genotyping.

Mouse DNA was isolated by adding lysing buffer (sodium hydroxide solution) to the samples for 2 hours at 55 °C, followed by neutralisation with a neutralisation buffer. DNA was captured using Promega MagneSil magnetic particles and immobilised with a magnet during three washes with ethyl alcohol. The DNA was eluted in nuclease-free water.

Primer mixes were prepared ahead of the experiment at 4× concentration and stored at -20 °C. When used, primers were diluted to 1.333×, yielding a final primer concentration of 900 nM. The 1.333× primer solution was combined with isolated DNA, and reactions were run on a ViiA 7 Real-Time PCR System under the following cycling conditions: 50 °C for 2 min and 95 °C for 10 min, followed by 40 cycles of 95 °C for 15 s and 60 °C for 1 min.

Primer sequences used for the genotyping are listed in the **Table 1**.

**Table 1.**
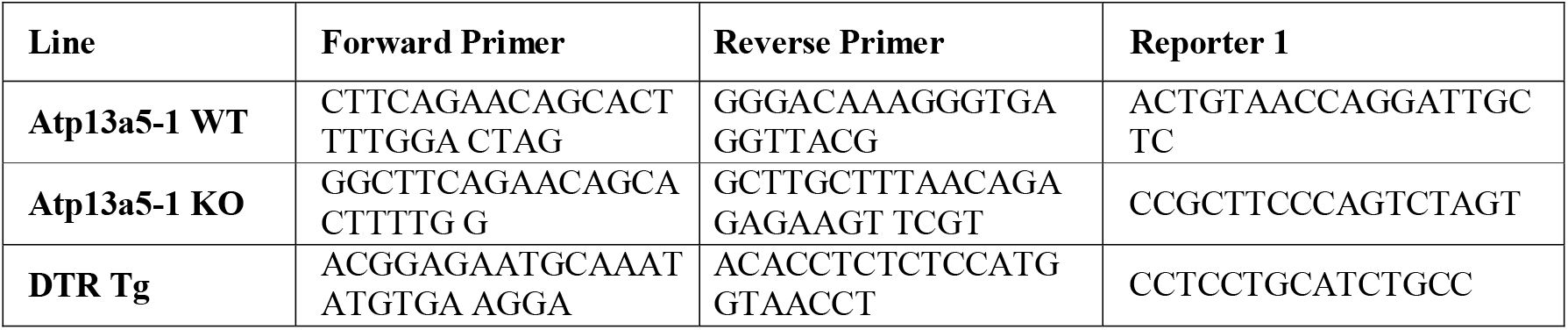
Summary of the primer sequences used by Transnetyx to genotype the *Atp13a5*^*tdT*^ and *Atp13a5;iDTR* mouse lines.

### Diphtheria toxin-dependent pericyte ablation

Homozygous *Atp13a5;iDTR* mice express tamoxifen-inducible DTR specifically in Atp13a5-positive cells. To ablate pericytes, tamoxifen (100 mg/kg; Sigma-Aldrich, T5648-5G) dissolved in corn oil (Sigma-Aldrich, 8001-30-7) was administered via oral gavage (100 μL/day) for five consecutive days. Tamoxifen was sonicated at 37 °C to aid dissolution and stored at 4 °C in the dark for the duration of the experiment.

Based on previous optimisation, DTR expression peaks 14 days after the final gavage. At this point, mice received intraperitoneal injections (i.p.) of diphtheria toxin (DT; Sigma-Aldrich, D0564-1MG) (1μg in 100 μL of sterile saline) or saline for five consecutive days.

Mice that lost >15% of their initial body weight or showed distress were humanely euthanised in accordance with license guidelines. Healthy mice were either sacrificed for tissue or imaged by MRI at days 5, 15, and 30 post-DT injection. These time points correspond to peak pericyte loss (day 5) and early and late recovery phases (days 15 and 30).

### Immunohistochemistry

Animals were euthanised by intraperitoneal injection of 100 μL pentobarbital. The anaesthetised mice were perfused transcardially with 15 mL of phosphate-buffered saline (PBS) containing 0.005 M ethylenediaminetetraacetic acid (EDTA) and post-fixed for 24 hours at 4 °C in 10% formalin solution. The following day, tissues were washed three times with PBS and incubated in 20% sucrose for 48 hours at 4 °C. Tissues were then flash-frozen in isopentane cooled with dry ice and stored at -80 °C until cryosectioning.

Brains were sectioned at 40 μm using a cryostat and stored at -20 °C in antifreeze solution (500 mL PBS (0.1 M sodium phosphate buffer (pH 7.4)), 85.6 g of sucrose (259 mM), 0.6 g of MgCl2 (7 mM), and 413.74 mL of glycerol).

For immunostaining, free-floating sections were washed and blocked for 2 hours at room temperature in 10% normal donkey serum (Bio-Rad Assay, bovine serum albumin standard), 0.1% Triton X-100 in 0.01 M PBS. Following blocking, 250 μL of primary antibodies (**Table 2**) were diluted in blocking solution and incubated overnight at 4 °C in the dark.

**Table 2.**
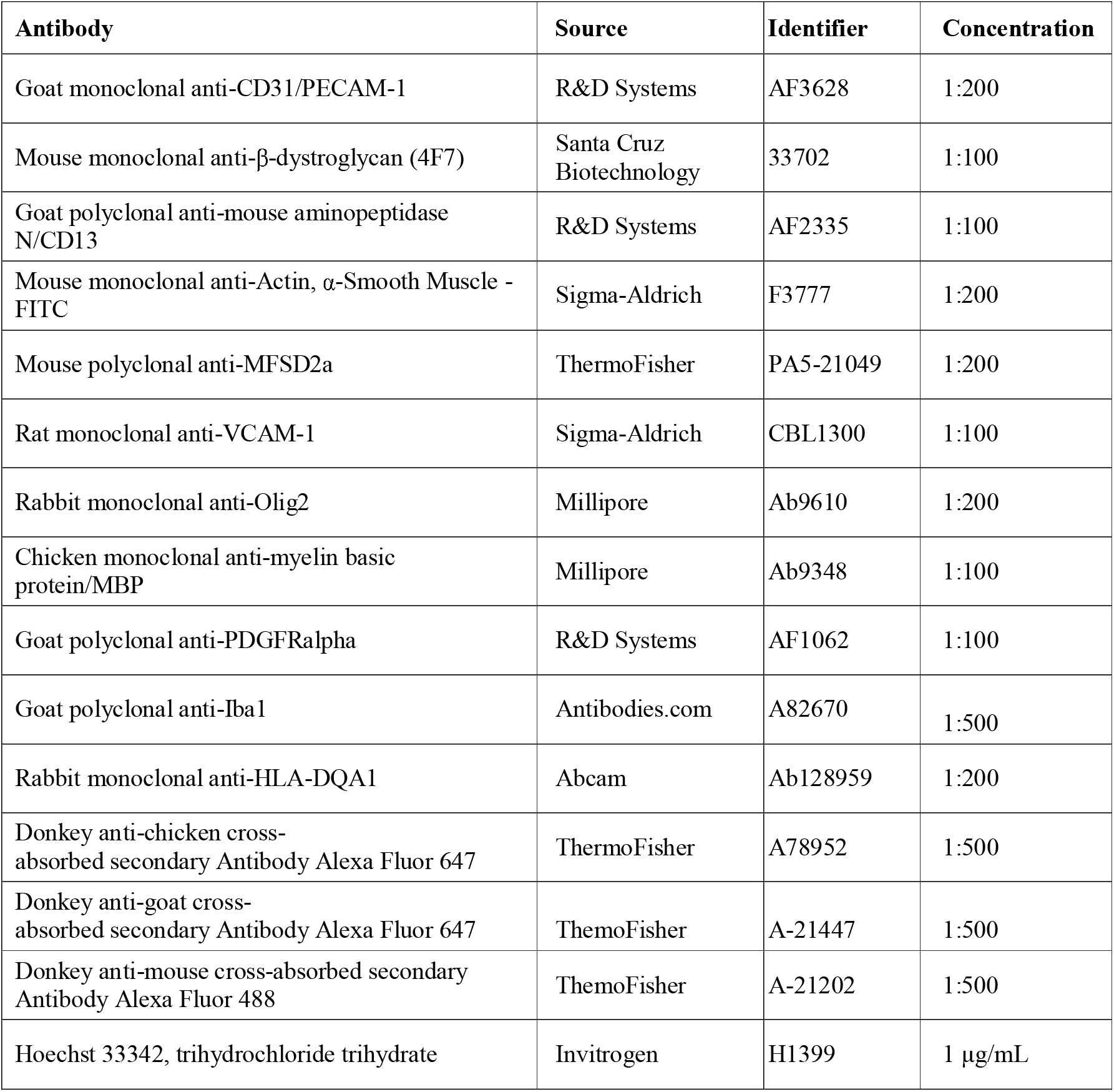
List of primary and secondary antibodies used in the study. The table includes name of antibody, source, identifier, and concentration used to achieve immunohistochemical staining of the mouse brain tissue.

| Antibody | Source | Identifier | Concentration |
| --- | --- | --- | --- |
| Goat monoclonal anti-CD31/PECAM-1 | R&D Systems | AF3628 | 1:200 |
| Mouse monoclonal anti-β-dystroglycan (4F7) | Santa Cruz Biotechnology | 33702 | 1:100 |
| Goat polyclonal anti-mouse aminopeptidase N/CD13 | R&D Systems | AF2335 | 1:100 |
| Mouse monoclonal anti-Actin, α-Smooth Muscle - FITC | Sigma-Aldrich | F3777 | 1:200 |
| Mouse polyclonal anti-MFSD2a | ThermoFisher | PA5-21049 | 1:200 |
| Rat monoclonal anti-VCAM-1 | Sigma-Aldrich | CBL1300 | 1:100 |
| Rabbit monoclonal anti-Olig2 | Millipore | Ab9610 | 1:200 |
| Chicken monoclonal anti-myelin basic protein/MBP | Millipore | Ab9348 | 1:100 |
| Goat polyclonal anti-PDGFRalpha | R&D Systems | AF1062 | 1:100 |
| Goat polyclonal anti-Iba1 | Antibodies.com | A82670 | 1:500 |
| Rabbit monoclonal anti-HLA-DQA1 | Abcam | Ab128959 | 1:200 |
| Donkey anti-chicken cross-absorbed secondary Antibody Alexa Fluor 647 | ThermoFisher | A78952 | 1:500 |
| Donkey anti-goat cross-absorbed secondary Antibody Alexa Fluor 647 | ThermoFisher | A-21447 | 1:500 |
| Donkey anti-mouse cross-absorbed secondary Antibody Alexa Fluor 488 | ThermoFisher | A-21202 | 1:500 |
| Hoechst 33342, trihydrochloride trihydrate | Invitrogen | H1399 | 1 μg/mL |

After overnight incubation, the primary antibody solution was discarded, and sections were washed three times for 10 minutes with 0.1% Triton-X in 0.01 M PBS. Conjugated secondary antibodies (**Table 2**), diluted in blocking solution, were then added and incubated for 2 hours at room temperature in the dark.

Following secondary antibody incubation, sections were washed once for five minutes with PBS. Nuclear staining was performed using Hoechst diluted in 0.1% Triton-X in 0.01 M PBS and incubated for five minutes at room temperature. Sections were then washed three times with PBS for 10 minutes each, mounted on slides using ProLong Gold antifade reagent, and coverslipped for imaging on the Opera Phenix confocal microscope.

For Iba1 and HLA-DQA1 immunostaining, 40 μm brain sections were processed using a modified antigen-retrieval protocol. Following fixation in 4% paraformaldehyde, sections were washed in PBS and antigen retrieval was performed in Tris-EDTA buffer containing 10 mM Tris base and 1 mM EDTA, pH 8.6, for 30 min at approximately 65 °C. Sections were then blocked and permeabilised in PBS containing 10% normal donkey serum and 0.2% Triton X-100 before overnight incubation at 4°C with goat anti-Iba1 and rabbit anti-HLA-DQA1 antibodies (**Table 2**).

As for standard immunohistochemistry, primary antibodies were discarded, and conjugated secondary antibodies diluted in blocking solution were added and incubated for 2 hours at room temperature in the dark. Sections were then washed in PBS, mounted on slides using ProLong Gold antifade reagent, and coverslipped.

### Image acquisition and processing

Mounted brain sections were imaged using an Opera Phenix spinning-disk confocal microscope. Whole brain images were acquired using a 20x water immersion objective (NA 1.0). Imaging channels included Hoechst, Alexa Fluor 488, endogenous tdTomato (Alexa Fluor 568), and Alexa Fluor 647.

Z-stack images were acquired across 30 planes with a 1-μm step size. Maximum-intensity projections were used for downstream analyses.

### IMARIS-based 3D rendering of pericyte morphotypes

*Atp13a5*^*tdT*^ brain sections immunostained for β-dystroglycan were used to acquire high-resolution representative images for three-dimensional rendering of pericyte morphotypes. Z-stacks spanning 20 μm were acquired from the cortex at Nyquist sampling using a Leica TCS SP8 confocal microscope equipped with a 40× oil-immersion objective (HC PL APO CS2, NA 1.30). Images were collected with a voxel size of 0.047 × 0.047 × 0.30 μm and deconvolved using Huygens Professional software (Scientific Volume Imaging, The Netherlands).

For 3D reconstruction, deconvolved image stacks were imported into IMARIS v.9.9 (Bitplane, Oxford Instruments). Pericytes labelled by endogenous tdTomato signal and vascular-associated β-dystroglycan signal were segmented independently using the Surface module.

For the β-dystroglycan channel, the surface grain size was set to 0.8 μm, background subtraction was performed using the diameter of the largest sphere set to 8.25 μm, and threshold were manually adjusted. For the pericyte channel, smoothing was enabled, the surface grain size was set to 0.095 μm, and background subtraction was performed using the diameter of the largest sphere set to 5.14 μm.

Background-associated surfaces were removed using volume filtering, generating a new channel containing surfaces larger than 10.0 μm^3^. Reconstructed 3D surfaces were visually inspected to confirm accurate rendering of pericyte morphology and vessel-associated β-dystroglycan signal before representative images were exported.

### Image Analysis

#### Pericyte coverage, vessel density, and vessel diameter

Pericyte coverage, vessel density, and vessel diameter were quantified using a custom FIJI (FIJI Is Just ImageJ, 10.1038/nmeth.2019) macro, VasOMatic (VOM)^17^.

Three 500 × 500 μm fields of view (FOV) per ROI were manually selected in the cortex, hippocampus, thalamus, and corpus callosum. Vascular segmentation was performed using VOM, which applied median smoothing, particle analysis, and manual thresholding to optimise vascular mask accuracy. The vascular mask served as a template for identifying tdTomato-positive Atp13a5 pericytes associated with blood vessels.

FIJI’s watershed function was then applied to divide the vasculature into discrete vascular segments, from which spatial metrics were extracted, including vessel length, diameter, pericyte coverage, and pericyte signal intensity, using skeleton measurements, distance mapping, and mask overlap approaches.

Pericyte coverage was expressed as the percentage of vessel area covered by pericytes. Vessel density was expressed as mm.mm^-3^, and vessel diameter in μm. Pericyte signal intensity was expressed as arbitrary units. Only vessels <10 μm in diameter were included, focusing analyses on capillaries and thin-strand pericytes. All segmentation thresholds were manually reviewed and adjusted to ensure segmentation accuracy.

#### Pericyte cell number

Pericyte cell numbers were quantified as densities to account for regional brain volume (cell count per volume) and local vascular density (cell count per vascular length), both of which can influence absolute counts. To estimate pericyte density in brain tissue, 2D projections of z-stacks from whole-brain confocal images were imported into QuPath, and 2-3 smaller ROIs per brain region were manually segmented. Each ROI encompassed all vascular beds and pericyte subtypes, owing to software limitations. These segmented ROIs were then exported and analysed using a custom MATLAB script.

Briefly, user-defined median filtering, adaptive thresholding based on the image mode, particle filtering, and smoothing were used to isolate tdTomato signal-rich pericyte somata. Centroids of identified somata were overlaid onto the original image for visual verification.

The volumes of ROIs were estimated based on slice thickness and FOV area. Pericyte density was expressed as the total number of identified somata divided by ROI volume (cells.mm^-3^). To normalise pericyte density relative to vascular density, vascular length was computed by the VOM script, and pericyte density values were divided by vascular density, yielding the number of pericytes per unit vessel length (cells.mm^-1^).

#### Spatial mapping and classification of pericyte subtypes

To assess spatial and subtype-specific changes in pericyte populations, we employed custom MATLAB-based tools developed in the laboratory. Pericytes somata were first identified and localised using MorphOMatic (MOM)^18^. Whole-brain fluorescence microscopy images were pre-processed by tiling (tile size = 500 pixels) to optimise memory usage.

Segmentation of the pericyte channel was achieved using a dual-mask flooding approach, whereby one mask identified pericyte somata and the second defined the immediate perivascular area. Upon completion of segmentation, MOM exported CSV files containing the X- and Y-coordinates of each soma, providing the spatial framework for subtype classification.

Pericyte subtype classification was performed using a ML-based convolutional neural network (CNN) implemented in MATLAB (version R2021b). The CNN architecture included an input layer of 128 × 128 × 1 and a classification output layer of 1 × 1 × 10.

The network was trained on 7,400 manually annotated images (128 × 128 pixels) of Atp13a5-positive pericytes, encompassing ten categories: background, lipofuscin, red blood cells, pericyte processes, vascular smooth muscle cells, ensheathing pericytes, mesh pericytes, junctional pericytes, *en passant* pericytes, and stellate pericytes. Junctional and *en passant* pericytes were grouped as thin-strand pericyte subtypes for downstream statistical analyses.

Eighty percent of images were used for training and 20% for testing, yielding a classification accuracy of 93%. For each pericyte soma detected by MOM, a corresponding 128 × 128 pixel snapshot of the Atp13a5 channel was extracted and classified by the CNN, enabling spatially resolved subtype mapping across the entire brain section.

#### Analysis of white matter integrity: myelin-based protein (MBP)

MBP staining was used as a one of the parameters to assess white matter integrity histologically. Myelin density was analysed using MOM^18^. Whole-brain fluorescence MBP images were pre-processed tile-wise on 1,000 x 1,000 pixels tiles at a resolution of 0.3 μm per pixel.

MBP staining was segmented using the Sobel edge-detection algorithm, and the 9,999/10,000^th^ quantile of the edge signal was selected as the thresholding value. Particles larger than 150 pixels were excluded, and the resulting mask was eroded by 1 pixel.

Following segmentation, MOM exported CSV files containing MBP-positive area measurements for each tile (in pixel units). The corpus callosum was manually delineated and applied to the exported dataset to extract ROI-specific data. Raw data were filtered to include only microvasculature-associated regions (vessel diameter 3-10 μm). The sum of MBP-positive tile areas was calculated, converted to μm^2^, normalised to corpus callosum ROI area, and expressed as percentage ROI area.

#### Analysis of white matter integrity: Olig2- and PDGFR_α_-positive cells

Olig2 and PDGFRα immunostaining were used to assess cellular correlates of white matter integrity. Whole coronal brain sections were analysed in QuPath, and the corpus callosum was manually delineated as the ROI for each section.

Cells were detected within each ROI using the Hoechst channel and QuPath watershed cell detection workflow. Detection parameters were optimised visually to identify Hoechst-positive nuclei while minimising background detection. Standard settings included a requested pixel size of 0.5 μm, background radius of 8 μm, sigma of 1.5 μm, minimum nuclear area of 10 μm^2^, maximum nuclear area of 150 μm^2^, and cell expansion of 3 μm.

Olig2 positivity was defined using nuclear Olig2 intensity, whereas PDGFRα positivity was defined using whole-cell PDGFRα intensity. Cells were classified in QuPath using scripted thresholds: Olig2-positive cells were defined as those with nucleus: Olig2 mean intensity ≥800, and PDGFRα-positive cells as those with cell: PDGFRα mean intensity ≥1,200.

Cells were classified as Olig2-only, PDGFRα-only, Olig2/PDGFRα double-positive, or negative. Marker-positive populations were expressed as a percentage of total Hoechst-positive cells.

Quality control was performed at both ROI and section level. ROIs with no detected cells, implausibly low cell density, missing or duplicate labels, or obvious detection failure were reviewed and reprocessed where appropriate.

#### Assessment of MFSD2A signal

Major Facilitator Superfamily Domain-containing protein 2A (MFSD2A) immunostaining was analysed using MOM to validate single-cell RNA-sequencing findings. Whole-brain fluorescence images were pre-processed tile-wise on 1,000 x 1,000 pixel tiles at a resolution of 0.3 μm per pixel.

MFSD2A signal was median-smoothed with a 5-pixel radius, thresholded using the mean intensity plus 0.5 × standard deviation, and median-smoothed again with a 5-pixel radius. Particles smaller than 200 pixels were removed, holes smaller than 500 pixels were filled, and a final median smoothing step with a 7-pixel radius was applied.

Individual vascular branches were segmented based on branches longer than 10 pixels from the mask skeleton.

Upon completion of segmentation, MOM exported CSV files containing MFSD2A signal measurements for each tile. The cortex was manually delineated and applied to exported dataset to restrict analyses to the ROI of interest. Average MFSD2A signal intensity was then calculated across tiles.

#### Assessment of VCAM-1 coverage

VCAM-1 immunostaining was analysed using MOM to assess endothelial activation following pericyte depletion. Whole-brain fluorescence images were pre-processed tile-wise on 1,000 × 1,000 pixel tiles at a resolution of 0.3 μm per pixel.

VCAM-1 signal was median-smoothed with a 5-pixel radius, thresholded using the mean intensity plus 0.5 × standard deviation, and median-smoothed again with a 5-pixel radius. Particles smaller than 200 pixels were removed, holes smaller than 500 pixels were filled, and a final median smoothing step with a 7-pixel radius was applied.

Individual vascular branches were segmented based on branches longer than 10 pixels from the mask skeleton. Upon completion of segmentation, MOM exported CSV files containing VCAM-1 coverage measurements for each tile.

The cortex was manually delineated and applied to the exported dataset to restrict analyses to the ROI of interest. VCAM-1 coverage was expressed as the percentage of vascular area positive for VCAM-1.

### Magnetic resonance imaging (MRI)

MRI was conducted at 5 (acute), 15 (progressive), and 30 (recovery) days post-DT injection on a 9.4 Tesla Bruker BioSpec system (BioSpec 94/20 USR, Bruker), equipped with 440 mT/m gradient strength. Each mouse (n = 8 per group) underwent two imaging sessions per time point under isoflurane anaesthesia in a 50:50 oxygen/air mixture at 1 L/min. Physiological parameters, including respiratory rate and core body temperature, were continuously monitored (SA Instruments). Respiration was maintained at 80-120 breaths/min, and body temperate at 36.5 ± 0.5 °C.

The first imaging session was acquired using an 86-mm transmit RF coil paired with a four-channel receiver-array coil (Bruker). Sequences included:

- T2-weighted rapid acquisition with relaxation enhancement (RARE) [TR/TE = 3,600/44 ms, RARE factor = 2, in-plane resolution 70 × 70 μm^2^, slice thickness 0.5 mm]
- Susceptibility-weighted imaging (SWI) [TR/TE = 662/8 ms, in-plane resolution 100 × 100 μm^2^, slice thickness 0.5 mm]
- Arterial spin labelling (ASL) [TR/TE = 10,000/10.91 ms, TI = 12 to 2,300 ms x 12, in-plane resolution 187.5 × 187.5 μm^2^, slice gap 1 mm]. Anaesthesia was reduced during ASL to minimise the vasodilatory effect of isoflurane^19^.
- Diffusion tensor imaging (DTI) [TR/TE = 2,000/16.9 ms, 30 diffusion directions, b-value = 670 s mm^-2^, in-plane resolution 167 × 167 μm^2^, slice thickness 0.74 mm].

The second session employed a 23-mm circularly polarised transmit-receive volume coil (Bruker). Sequences included:

- T2-weighted [same parameters as above except RARE factor = 6]
- T1-weighted FLASH [TR/TE = 19.19/3.8 ms, flip angles = 5°, 10°, 15°, 30°, and 45°, in-plane resolution 159 × 159 μm^2^, slice thickness 1 mm, slice gap 1 mm, two averages]
- Dynamic contrast-enhanced (DCE) [TR/TE = 19.19/3.8 ms, flip angle = 15°, temporal resolution = 4.34 s, 332 volumes]. Gadolinium (Gd-DTPA; BioPAL, Inc.; 1:15 dilution in 1% EDTA-saline) was administered via tail vein after 4 minutes at a rate of 600 μL.min^-1^ using a custom infusion system.

### MRI post-processing analysis

All MRI datasets were converted to DICOM format using Paravision 360 version 3.3 (Bruker) and subsequently converted to nifti using *BrkRaw* prior to downstream analyses.

#### Cerebral blood flow analysis

ASL-derived CBF maps were generated using Paravision 360 (Bruker). CBF values (in mL.min^-1^.100g^-1^) were extracted from manually delineated ROIs drawn on T1-weighted images in FIJI/ImageJ.

ROIs included anterior and posteriorly cortex, corpus callosum, caudate putamen, hippocampus, thalamus, and internal capsule.

#### Blood-brain barrier permeability analysis

DCE-MRI datasets were processed the ROCKETSHIP MATLAB toolbox^20^. BBB permeability (blood-to-brain transfer constant, K_trans_) was quantified using Patlak modelling for the same ROIs used in CBF analyses.

The tissue contrast concentration Ct_issue_(t) was modelled as a function of the vascular input function (VIF), plasma volume fraction (v_p_), and K_trans_^21^.

VIFs were automatically extracted using ROCKETSHIP build-in functions by selecting physiologically consistent voxels across all mice.

### Diffusion tensor imaging analysis

Diffusion tensor fitting was performed using the FMRIB Software library (FSL) Diffusion Toolbox (FDT). Non-linear registration to the MouseX DW-Allen Atlas^22^ was performed using Advanced Normalization Tools (ANTs, version 2.6.2)^23^.

Atlas and template brain masks were warped back into individual subject space for skull stripping and extraction of scalar diffusion metrics, including fractional anisotropy (FA), mean diffusivity, axial diffusivity, and radial diffusivity.

A white matter skeleton (threshold = 0.2) and white matter mask (threshold = 0.25) were generated from the MouseX FA template image using FSL Tract-Based Spatial Statistics (TBSS). These were used for 500 voxel-wise permutations with Randomise (FSL), applying Threshold-Free Cluster Enhancement (TFCE), family-wise error multiple comparisons corrections, and a general linear model with a 2 x 2 design.

### Single-cell RNA sequencing: vessel isolation and library preparation

*Atp13a5;iDTR* mice treated with tamoxifen followed by diphtheria toxin (DT) or saline (CT) were transcardially perfused 5 or 30 days after the final DT injection (n = 4 per group). The olfactory bulbs, brainstem, and cerebellum were removed, and the remaining brain tissue was cut sagittally into eight pieces.

Tissue pieces were transferred to a C-tube (Miltenyi Biotec, 5221003005) containing 1,950 μL of enzyme mix 1 prepared from 50 μL Enzyme P and 1,900 μL Buffer Z, together with 30 μL of enzyme mix 2 prepared from 10 μL Enzyme A and 20 μL Buffer Y. All dissociation reagents were supplied as part of the Adult Brain Dissociation Kit (Miltenyi Biotec, 5250104875).

Tissue was dissociated for 30 min using the gentleMACS Octo Dissociator with Heaters. The resulting suspension was passed through a MACS SmartStrainer (Miltenyi Biotec, 9220800143) pre-wetted with 4 mL D-PBS (Gibco, 2886285), followed by a 10 mL D-PBS wash. The cell suspension was gently triturated and centrifuged at 300 × g for 10 min at 4 °C.

The pellet was resuspended in 3,100 μL cold D-PBS and transferred to a 15 mL tube containing 900 μL Debris Removal Solution. The suspension was mixed by pipetting ten times, and 4 mL D-PBS was carefully layered at a 20° angle to generate a density gradient. Samples were centrifuged at 3,000 × g for 10 min at 4 °C, generating three distinct layers consisting of debris, myelin, and target cells. The upper debris and myelin fractions were removed, and the remaining cell-containing fraction was centrifuged at 100 × g for 10 min at 4 °C.

Residual red blood cells were removed using Red Blood Cell Removal Solution for 10 min at 4 °C. Samples were centrifuged again at 300 × g for 10 min at 4 °C and resuspended in 550 μL PB buffer prepared as a 1:20 dilution of MACS Bovine Serum Albumin Stock Solution (Miltenyi Biotec, 130-091-376) in D-PBS.

Following removal of myelin, debris, and red blood cells, the pellet was resuspended in 80 μL PB buffer and incubated with 10 μL FcR Blocking Reagent (Miltenyi Biotec, 5240403597) for 10 min at 4 °C. CD31-positive vascular cells were labelled by addition of 10 μL CD31 MicroBeads (Miltenyi Biotec, 130-097-418) and incubation for 15 min at 4 °C in the dark.

To enrich the vascular fraction, the suspension was passed through pre-wetted pre-separation filters (Miltenyi Biotec, 9221100219) attached to magnetic columns attached to an OctoMACS separator. The CD31-positive fraction was centrifuged at 300 × g for 10 min at 4 °C, and the resulting pellet containing brain microvessels and associated cells was resuspended in PB buffer.

RNA was isolated from the vascular fraction, and single-cell libraries were prepared using the 10x Genomics Chromium Single Cell 3′ v4 kit according to the manufacturer’s instructions, including cell capture, barcoding, reverse transcription, and cDNA amplification. Library quality was assessed using the Agilent TapeStation HS D1000 assay.

Sequencing was performed on an Illumina NovaSeq platform using four 100-cycle runs on a P4 flow cell. Sequencing depth ranged from 372.5 to 532.3 million paired-end reads per library, with a mean depth of 456.1 million paired-end reads.

### Single-cell RNA sequencing: data analysis

#### Quality control and cluster identification

Raw sequencing data generated on the Illumina NovaSeq platform were demultiplexed and processed using Cell Ranger v6.0.1 (10x Genomics). Reads were aligned to the mouse reference genome mm10/GRCm38, and gene-cell count matrices were generated for downstream analyses. Gene count matrices for each individual sample were imported into Seurat (v4.3).

For comparison with published cerebrovascular scRNA-seq datasets (**Fig. 1C**)^13^, data were downloaded from the Gene Expression Omnibus under accession GSE98816 and re-processed in Seurat v4.3 using the same downstream analysis framework. For the analyses shown in Figure 1, 1,966 cells from the present dataset and 1,331 cells from the Vanlandewijck et al. dataset were included.

**Figure 1.**
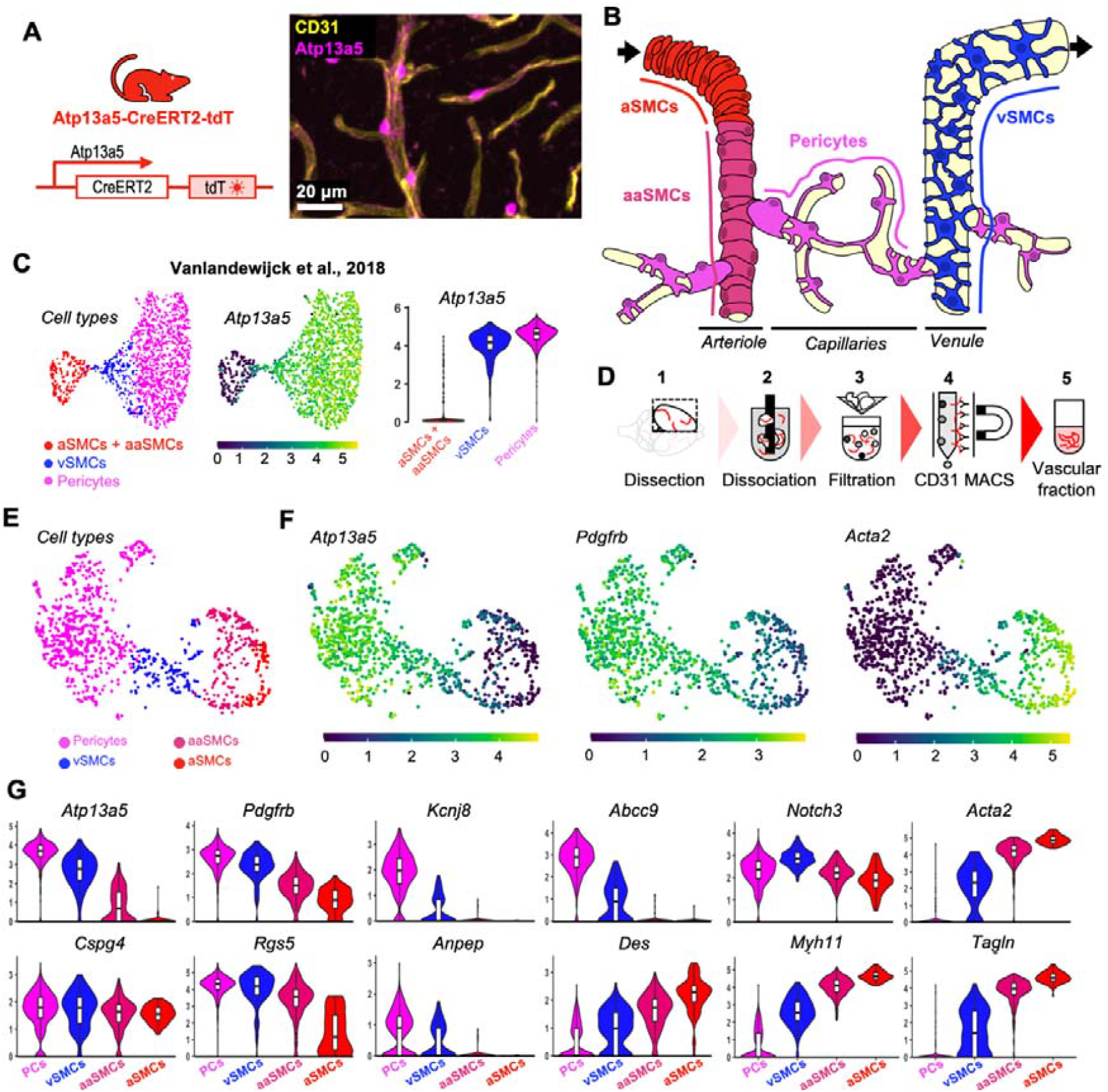
Atp13a5 selectively labels pericytes and defines a mural cell transcriptional signature. (**A**) Schematic of the genetic strategy used to generate the *Atp13a5*-CreERT2-tdTomato reporter line. Representative confocal image showing tdTomato-positive pericytes closely associated with CD31-positive vessels. Scale bar: 20 μm. (**B**) Schematic representation of the distribution of mural cell subtypes along the arterio-venous axis, including arterial smooth muscle cells (aSMCs), arteriole-associated smooth muscle cells (aaSMCs), capillary pericytes, and venous smooth muscle cells (vSMCs). (**C**) Single-cell analysis of brain mural cells from a publicly available single-cell RNA-sequencing (scRNA-seq) dataset^13^, showing mural cell subtype distribution, *Atp13a5* expression, and relative abundance of each subtype. (**D**) Workflow for scRNA-seq of vascular cells isolated from mouse brain tissue using CD31-based magnetic-activated cell sorting (MACS). (**E**) UMAP showing the distribution of mural cell subtypes, including pericytes, vSMC, aaSMC, and aSMC populations. (**F**) UMAP feature plots showing *Atp13a5, Pdgfrb*, and *Acta2* expression across mural cell subtypes in the scRNA-seq dataset generated in this study (n = 6 mice). (**G**) Violin plots showing quantitative expression levels of canonical mural cell markers across the four identified clusters.

Doublets identified by *scDblFinder*^24^ and cells containing >10% mitochondrial transcripts were excluded, resulting in 169,204 cells retained for downstream analyses.

Datasets were normalised with Seurat’s *NormalizeData()* function, and the top 2,000 variable genes were identified using *FindVariableFeatures()*. Principal component analysis (PCA) and elbow plot inspection were used to determine the optimal number of principal components for clustering.

Clusters were identified using Seurat’s *FindClusters()* function across multiple resolutions (0.1-1.5), and the optimal clustering resolution was selected using the *clustree* package^25^. Clusters were annotated based on expression of well-established canonical marker genes.

For subclustering analyses, annotated cell populations of interest were extracted, and the clustering workflow was repeated following re-identification of the top 2,000 variable genes.

#### Differential expression, enrichment, and pathway analyses

Differential expression analysis for each cell type was performed using the *MAST R* package^26^. Following exclusion of lncRNA genes and log-normalisation of raw counts, two-part hurdle models were fitted to identify differentially expressed genes (DEGs), with treatment group, gene count, mitochondrial and ribosomal percentages included as fixed effects, and sample ID included as a random effect.

Genes with Benjamini-Hochberg adjusted p-value < 0.05 were considered as DEGs.

DEG lists for each comparison were ranked according to z-score and analysed using *clusterProfiler* for gene-set enrichment analysis (GSEA) against the KEGG database and lists of GWAS-nominated genes associated with AD and SVD, including WMH and PVS datasets^12^.

Transcription factor activity was inferred using the *decoupleR* package^27^ using the same ranked gene lists used for GSEA analyses.

### Statistical analysis

All statistical analyses were performed in R and RStudio. Linear mixed-effects models (*lmerTest* package) were used to assess statistical significance across datasets, incorporating fixed effects (Treatment, Timepoint, ROI, and Sex) and random effects (Mouse ID) to account for repeated measures while maximising statistical power.

Model assumptions, including linearity, homogeneity of variance, and normality, were assessed using the *check_model()* function from the *performance* package. For BBB permeability measurements and VCAM-1 coverage, data was inverse-normal transformed where required to satisfy model assumptions.

For datasets with multiple continuous dependent variables, data were standardised using the *scale()* function in R.

Post hoc comparisons were performed using the *estimate_contrasts()* function from the *modelbased* package, which computes estimated marginal means, confidence intervals, and p-values for pairwise comparison of interest. P-values were adjusted for multiple testing using Bonferroni, false discovery rate (FDR), or Tukey correction, as appropriate. Statistical significance was defined as follows: *p* < 0.05 = *, *p* < 0.01 = **, *p* < 0.001 = ***, 0.05 ≤ *p* < 0.1 = # (trend), n.s. = not significant.

## Results

### Atp13a5 selectively labels brain pericytes and defines a mural cell transcriptional program

To assess the specificity of the *Atp13a5* mouse line for targeting brain pericytes, we used the inducible *Atp13a5*-CreERT2-tdTomato reporter line^16^ (**Fig. 1A**). Confocal imaging revealed tdTomato-positive cells closely associated with CD31-positive vessels, displaying morphology consistent with perivascular mural cells (**Fig. 1A**). To position Atp13a5-expressing cells across the vascular tree, we considered the distribution of mural cell subtypes, including arterial smooth muscle cells (aSMCs), arteriole-associated smooth muscle cells (aaSMCs), capillary pericytes, and venous smooth muscle cells (vSMCs) (**Fig. 1B**). Re-analysis of a publicly available scRNA-seq dataset^13^ demonstrated that Atp13a5 expression is enriched in pericytes compared to other mural cell subtypes (1,331 cells; **Fig. 1C**). We next performed scRNA-seq on vascular-enriched brain fractions isolated using CD31-based MACS (**Fig. 1D**) and also identified distinct mural cell populations (1,966 cells) corresponding to pericytes, vSMCs, aaSMCs, and aSMCs (**Fig. 1E**). Feature plots showed that *Atp13a5* expression co-localised with canonical pericyte markers such as *Pdgfrb*, while being largely absent from *Acta2*-positive SMC populations (**Fig. 1F**). Quantitative analysis of marker expression confirmed that pericyte clusters were enriched for *Atp13a5, Pdgfrb, Kcnj8*, and *Abcc9*, whereas SMC clusters preferentially expressed *Acta2, Des, Myh11*, and *Tagln* (**Fig. 1G**).

Together, these data demonstrate that Atp13a5 selectively labels a transcriptionally defined pericyte population within the brain microvasculature.

### Pericyte subtypes display distinct morphologies and spatial distributions across the vascular tree

We next characterised pericyte subtypes at the tissue level based on morphology and vascular localisation. Three distinct morphotypes - ensheathing (ES), mesh (M), and thin-strand (TS) pericytes - were identified along the vascular hierarchy (**Fig. 2A,B**). Multiplex immunofluorescence confirmed that these morphotypes could be distinguished based on their relationship to α-SMA-positive vessels and CD13-positive perivascular localisation (**Fig. 2C,D**). tdTomato expression in CD13-positive cells was observed across multiple brain regions and along the vascular branching hierarchy, spanning precapillary arterioles (PCA), capillaries, postcapillary venules (PCV), and ascending venules (AV) (**Fig. 2E**). Quantification revealed that Atp13a5/CD13 double-positive pericyte coverage varied as a function of vessel diameter (**Fig. 2F**), consistent with subtype-specific distribution.

**Figure 2.**
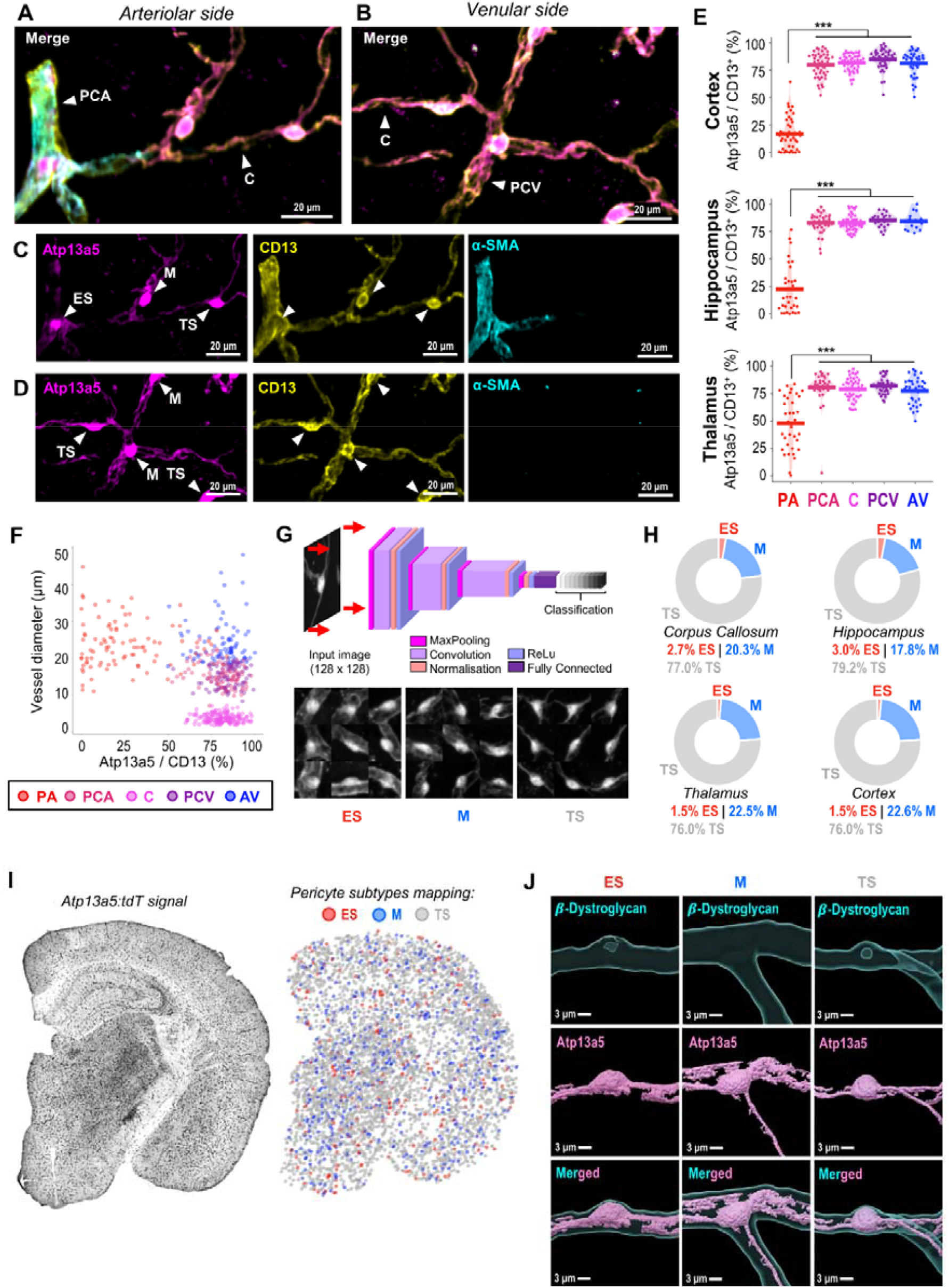
Morphological and spatial characterisation of pericyte subtypes across the vascular tree. (**A**) Ensheathing, mesh, and thin-strand pericytes distributed along increasing branching orders of precapillary arterioles (PCA) and capillaries (C). Scale bar: 20 μm. (**B**) Mesh and thin-strand pericytes located on capillaries (C) transitioning into postcapillary venules (PCV). Scale bar: 20 μm. (**C, D**) Multiplex immunofluorescence images of Atp13a5, CD13, and α-SMA channels, indicating the positions of ensheathing (ES), mesh (M), and thin-strand (TS) pericytes. Scale bars: 20 μm. (**E**) tdTomato (tdT) expression in CD13+ perivascular cells across brain regions (cortex, hippocampus, and thalamus) and along the vascular branching hierarchy, including penetrating arterioles (PA), precapillary arterioles (PCA), capillaries (C), postcapillary venules (PCV), and ascending venules (AV). (**F**) Distribution of Atp13a5+/CD13+ pericyte coverage as a function of vascular diameter. (**G**) Architecture of the convolutional neural network (CNN) used to classify pericyte subtypes, along with representative 3 × 3 image examples used for training. (**H**) Proportion of pericyte subtypes in corpus callosum, hippocampus, thalamus, and cortex in baseline animals. (**I**) Spatial mapping of pericyte subtypes (ES: ensheathing, M: mesh, and TS: thin-strand) on a representative brain section from a baseline animal. (**J**) Three-dimensional rendering of representative ES, M, and TS pericytes, together with the vascular basement membrane (β-dystroglycan). Scale bars: 3 μm. Panels E-I: n = 6 biologically independent mice (3 males, 3 females). Statistical analyses were performed using linear mixed-effects models followed by Tukey’s post hoc multiple-comparisons correction (***, p < 0.001 - with treatment, timepoint, and sex as fixed effects and mouse ID as a random effect).

To enable unbiased classification, we developed a ML-based convolutional neural network (CNN) trained on thousands of annotated pericyte images (**Fig. 2G**). This approach enabled automated quantification of pericyte subtypes across brain regions. Analysis revealed region-specific distributions of ES, M, and TS pericytes (**Fig. 2H**), which were further visualised through spatial mapping across brain sections (**Fig. 2I**) and confirmed by three-dimensional reconstructions (**Fig. 2J**).

These data establish that pericyte subtypes exhibit distinct morphological and spatial organisation across the brain vascular tree.

### Inducible pericyte ablation leads to transient loss and subtype-specific remodelling

To investigate the functional role of Atp13a5-positive pericytes, we generated *Atp13a5;iDTR* mice enabling inducible ablation following diphtheria toxin (DT) administration (**Fig. 3A,B**). Pericyte depletion resulted in a marked reduction in pericyte coverage at 5 days post-treatment, which partially recovered by 30 days (**Fig. 3C,D**), corresponding to an average ∼10-15% reduction at 5 days, with region-specific decreases of 13.6% in the corpus callosum and 14.6% in the hippocampus, and recovery to near-control levels by 30 days. Similarly, pericyte somata density per vascular length was significantly reduced at 5 days, with a more pronounced decrease compared to coverage (-26.7% in the corpus callosum and -31.2% in the hippocampus), and returned towards baseline at 30 days (**Fig. 3E**). This differential effect between coverage and cell density is consistent with compensatory extension of processes from neighbouring pericytes.

**Figure 3.**
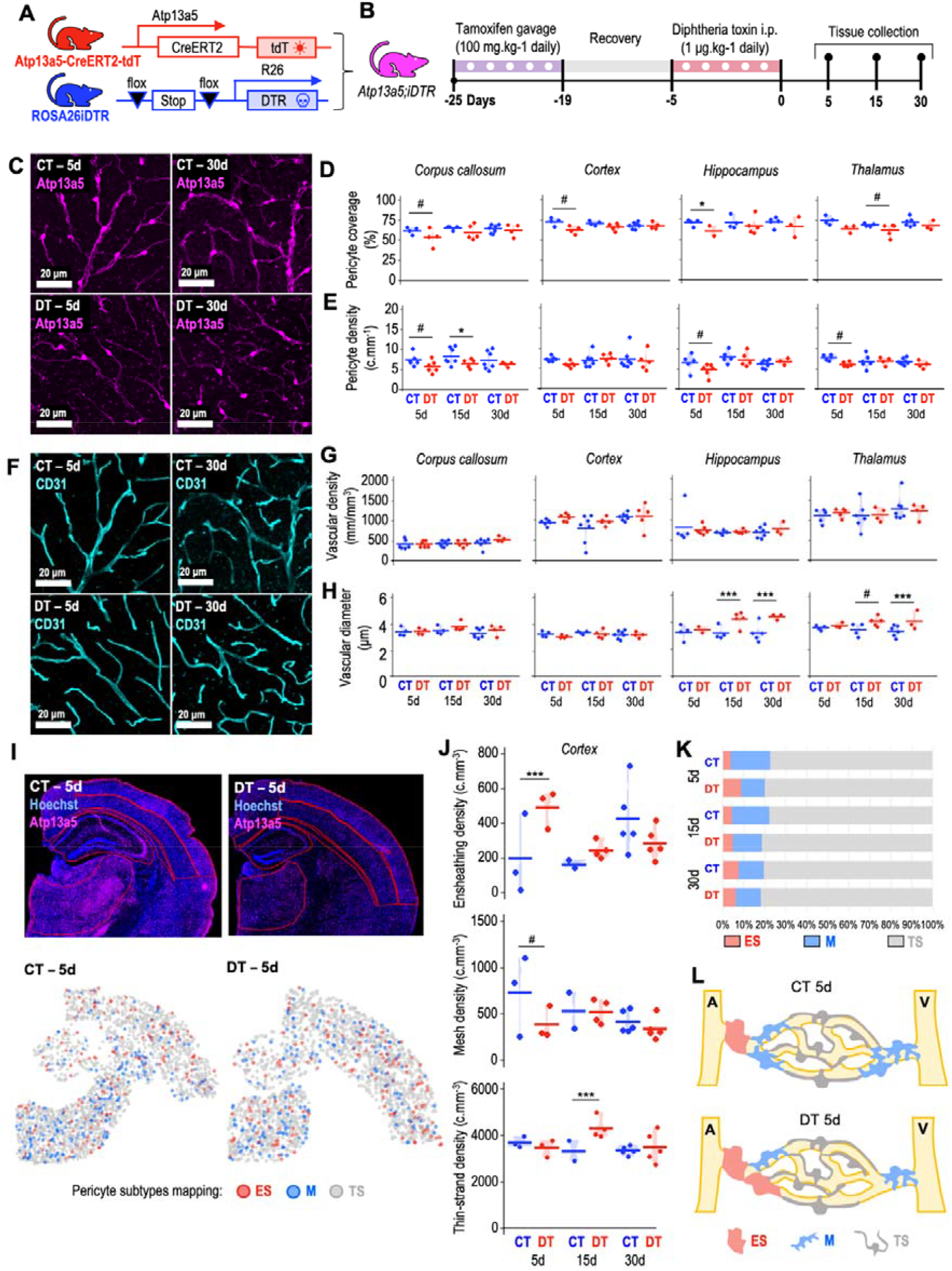
Inducible pericyte ablation leads to transient loss and subtype-specific remodelling. (**A**) Breeding strategy used to generate *Atp13a5;iDTR* mice. (**B**) Experimental timeline for pericyte depletion using diphtheria toxin (DT). (**C**) Representative images showing pericyte loss at 5- and 30-day post-depletion in DT-treated and control (CT) animals. Scale bars: 20 μm. (**D**) Quantification of pericyte coverage following depletion (n = 2-7 mice per group). (**E**) Pericyte somata density per vascular length following depletion (n = 4-7 mice per group). (**F**) Representative images of endothelial cell morphology (CD31 staining) at 5- and 30-day post-depletion in DT-treated and CT animals (n = 4-7 mice per group). Scale bars: 20 μm. (**G**) Vascular density following pericyte depletion (n = 3-7 mice per group). (**H**) Vascular diameter following pericyte depletion (n = 2-7 mice per group). (**I**) Regions selected for automated pericyte subtype identification using CNN, showing localisation of thin-strand (TS, grey), ensheathing (ES, red), and mesh (M, blue) pericytes in CT and DT-treated animals at 5 days post-depletion. (**J**) Quantification of pericyte subtype densities in the cortex (n = 2-5 mice per group). (**K**) Relative proportions of pericyte subtypes following depletion. (**L**) Summary schematic illustrating subtype-specific vulnerability to acute depletion. A: arteriole; V: venule; iDTR: inducible diphtheria toxin receptor; DT: diphtheria toxin; CT: control; i.p.: intraperitoneal; tdT: tdTomato; ES: ensheathing pericytes; M: mesh pericytes; TS: thin-strand pericytes. Statistical analyses were performed using linear mixed-effects models with FDR correction (***, p < 0.001; *, p < 0.05; ^#^, p = 0.05-0.1 - with treatment, timepoint, and sex as fixed effects and mouse ID as a random effect).

Despite pericyte loss, endothelial morphology appeared grossly preserved (**Fig. 3F**), and no significant changes were observed in vascular density (**Fig. 3G**). Vascular diameter was largely unchanged overall; however, region-specific increases were observed in the hippocampus and thalamus at later timepoints, with increases of 41.9% and 28.9% in the hippocampus, and 15% and 23.3% in the thalamus at 15- and 30-day post-depletion, respectively (**Fig. 3H**).

Using CNN-based classification, we next assessed pericyte subtype vulnerability. While all subtypes were affected, mesh pericytes showed a more pronounced reduction (-12.1%) compared to other subtypes (**Fig. 3J,K**). This resulted in a shift in subtype composition, with a relative increase in the proportion of ensheathing pericytes (+23.5%), despite an overall reduction in total pericyte numbers (**Fig. 3K,L**).

These findings indicate that acute pericyte ablation induces transient loss accompanied by subtype-specific remodelling of the pericyte population, with near-complete recovery of both coverage and cell density by 30 days post-depletion.

### Pericyte loss induces transient cerebrovascular dysfunction and white matter alterations

We next assessed the functional consequences of pericyte depletion using longitudinal MRI (**Fig. 4A**). ASL-derived CBF maps revealed a reduction in perfusion at 5 days post-depletion (**Fig. 4B,C**), with regional heterogeneity across brain regions but an overall reduction in brain perfusion (-21.7%), reaching significance in the cortex. CBF progressively recovered over time and returned to baseline levels by 30 days, paralleling the repopulation of pericytes observed in Fig. 3.

**Figure 4.**
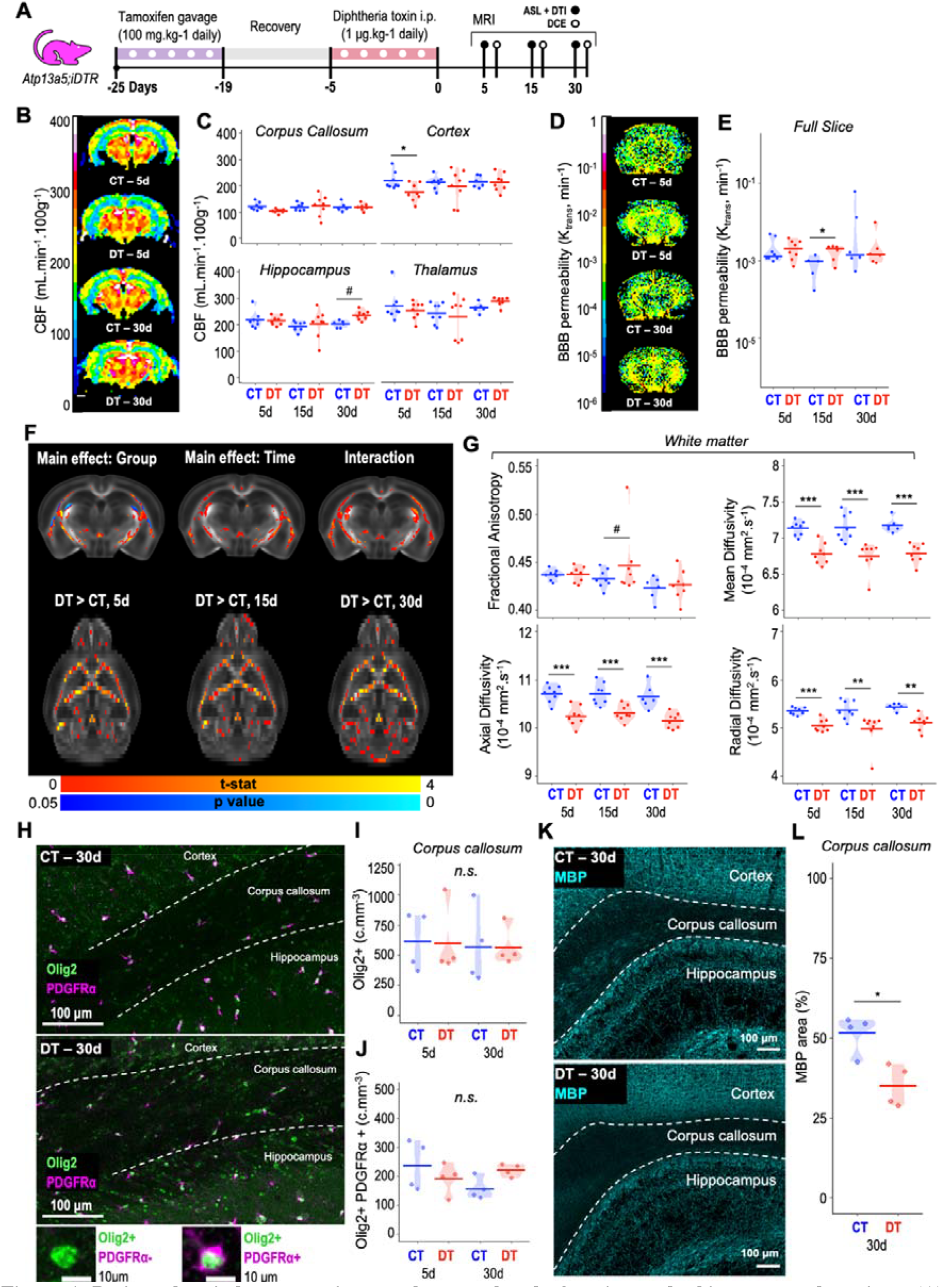
Pericyte loss induces transient cerebrovascular dysfunction and white matter alterations. (**A**) Experimental timeline of MRI acquisitions, including arterial spin labelling (ASL), dynamic contrast-enhanced (DCE), and diffusion tensor imaging (DTI). (**B**) Representative ASL-derived cerebral blood flow (CBF) maps at 5- and 30-day post-DT in pericyte-depleted (DT) and control (CT) mice. (**C**) Quantification of CBF changes over time in the corpus callosum, cortex, hippocampus, and thalamus (n = 6-8 mice per group). (**D**) Representative DCE-derived maps of blood-brain barrier (BBB) permeability (K_trans_) at 5- and 30-day post-DT in pericyte-depleted (DT) and control (CT) mice. (**E**) Quantification of whole brain BBB permeability changes over time (n = 5-8 mice per group). (**F**) Diffusion tensor imaging (DTI) metrics mapped onto a normalized template (MouseX), showing group- and time-dependent changes. (**G**) Quantification of DTI metric changes over time in white matter (n = 6-8 mice per group). (**H**) Representative images of Olig2/PDGFRα double-positive cells in the corpus callosum at 30-day post-depletion. Scale bars: 100 μm. (**I**) Olig2-positive cell density in the corpus callosum at days 5 and 30 post-depletion (n = 4 mice per group). (**J**) Olig2/PDGFRα double-positive cell density in the corpus callosum at days 5 and 30 post-depletion (n = 4 mice per group). (**K**) Representative images of MBP staining in the corpus callosum at 30-day post-depletion. (**L**) Quantification of MBP-positive area (%) in the corpus callosum at day 30 post-depletion (n = 4 mice per group). Scale bars: 100 μm. MRI: magnetic resonance imaging; ASL: arterial spin labelling; DCE: dynamic contrast-enhanced; DTI: diffusion tensor imaging; DT: diphtheria toxin; CT: control; BBB: blood-brain barrier; K_trans_: blood-to-brain transfer constant; PDGFRα: platelet-derived growth factor receptor alpha; MBP: myelin basic protein; Olig2: oligodendrocyte transcription factor 2. Statistical analyses were performed using linear mixed-effects models with FDR or Bonferroni correction (***, p < 0.001; **, p < 0.01; *, p < 0.05; ^#^, p = 0.05-0.1; n.s., not significant-with treatment, timepoint, and sex as fixed effects and mouse ID as a random effect).

DCE-MRI analysis showed mild but measurable increases in BBB permeability, with global BBB disruption reaching significance at day 15 post-depletion (**Fig. 4D,E**), corresponding to a 28.3% increase in K_trans_, which normalised towards control levels by 30 days.

DTI-MRI revealed mild but significant alterations in white matter microstructure, with decreases in mean, axial, and radial diffusivities across time (∼6% reduction) (**Fig. 4F,G**). At the cellular level, immunohistochemistry demonstrated no significant changes in Olig2-positive oligodendrocyte lineage cell density at 5- and 30-day post-depletion in the corpus callosum (**Fig. 4I**), but a trend towards an increased number of Olig2/PDGFRα double-positive oligodendrocyte progenitor cells (OPCs) at day 30 in pericyte-depleted mice compared to controls (**Fig. 4J**). At 30 days post-depletion, myelin basic protein (MBP) staining revealed a marked reduction in myelin content in the corpus callosum (∼40% decrease) (**Fig. 4K,L**).

Together, these data show that pericyte loss induces transient cerebrovascular dysfunction, characterised by reduced perfusion and increased BBB permeability, alongside white matter alterations associated with changes in oligodendrocyte lineage dynamics and myelin loss.

### Pericyte depletion induces capillary endothelial identity shift, interferon signalling, and enrichment of human SVD-associated gene signatures

To investigate downstream molecular effects, we performed scRNA-seq of CD31-enriched (MACS-based pull-down) vascular brain samples following pericyte depletion. Clustering identified multiple vascular and glial cell types, with 92% of captured cells corresponding to endothelial cells (ECs), alongside smaller populations of pericytes, smooth muscle cells, and immune cells (**Fig. 5A**). Subclustering of ECs resolved arterial (aECs), capillary (cECs), and venous (vECs) populations along the arteriovenous axis (**Fig. 5B**), defined by canonical markers including *Bmx*/*Efnb2* (arterial), *Mfsd2a* (capillary), and *Nr2f2*/*Aplnr* (venous).

**Figure 5.**
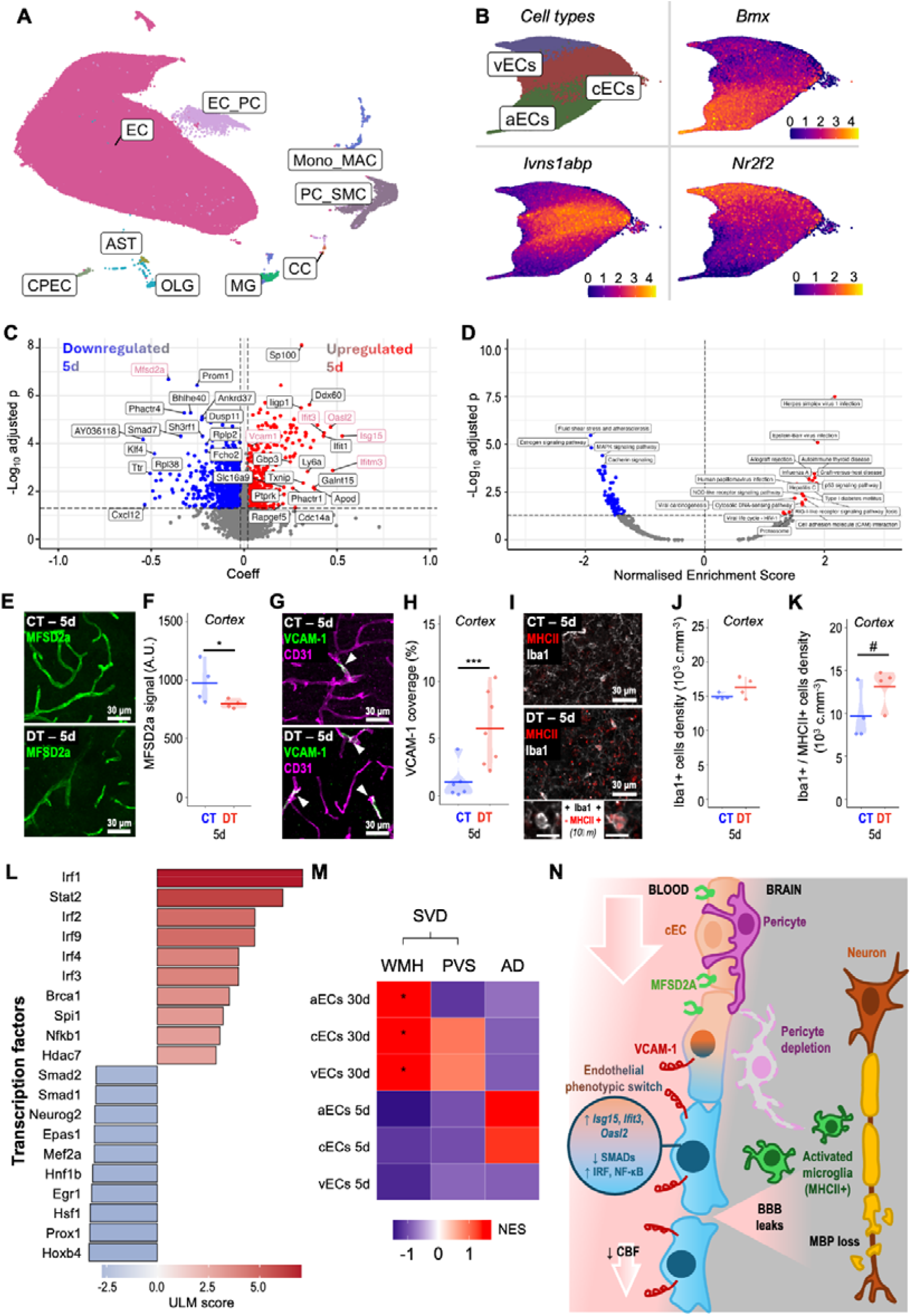
Acute pericyte depletion alters capillary endothelial identity, activates interferon signalling, and associates with human SVD gene signatures. (**A**) UMAP representation of cell clusters identified by scRNA-seq of vascular-enriched brain samples at 5 and 30 days post-depletion (169,204 cells), including endothelial cells (EC), pericytes (PC), smooth muscle cells (SMC), monocytes (Mono), macrophages (MAC), microglia (MG), astrocytes (AST), oligodendrocytes (OLG), choroid plexus epithelial cells (CPEC), and chondrocyte-like cells (CC). (**B**) UMAP of subclustered EC subtypes along the arteriovenous axis (155,363 cells), identifying arterial endothelial cells (aECs), capillary endothelial cells (cECs), and venous endothelial cells (vECs), distinguished by canonical markers including *Bmx*/*Efnb2* (arterial), *Mfsd2a* (capillary), and *Nr2f2*/*Aplnr* (venous). (**C**) Volcano plot of differentially expressed genes in cECs at 5 days post-depletion, with key genes of interest highlighted in pink, including markers of endothelial activation (e.g., *Vcam1*), interferon signalling (e.g., *Isg15, Ifit3*), and BBB function (e.g., *Mfsd2a*). Differential expression was performed using MAST (significantly upregulated: coef > 0.03, p_adj_ < 0.05; downregulated: coef < 0.03, p_adj_ < 0.05). (**D**) Volcano plot of KEGG pathways identified by GSEA, showing positively (normalised enrichment scores (NES) > 1.5) and negatively (NES < 1.5) enriched pathways (p_adj_ < 0.05) in cECs at 5 days post-depletion. (**E**) Representative images of MFSD2A expression in the cortex at 5-day post-DT in pericyte-depleted (DT) and control (CT) mice. (**F**) Quantification showing loss of MFSD2A signal in the cortex at 5 days post-depletion (n = 4 mice per group). (**G**) Representative images of VCAM-1 expression in the cortex at 5-day post-DT in pericyte-depleted (DT) and CT mice. (**H**) Quantification showing increased VCAM-1 coverage (endothelial activation) in the cortex at 5 days post-depletion (n = 6-7 mice per group). (**I**) Representative images of Iba1/MHCII double-positive microglia in the cortex at 5-day post-DT in pericyte-depleted (DT) and CT mice. (**J**) Iba1-positive cell density in the cortex at 5 days post-depletion (n = 4 mice per group). (**K**) Increase in Iba1/MHCII double-positive cell density in the cortex at 5 days post-depletion (n = 4 mice per group). (**L**) Waterfall plot showing top transcription factors with predicted increased (red) or decreased (blue) activity in cECs at 5 days post-depletion (decoupleR analysis). (**M**) Heatmap of GSEA NES against AD-, SVD WMH-, and SVD PVS-GWAS gene sets across EC subtypes at 5- and 30-day post-depletion, highlighting selective enrichment of SVD-associated signatures in endothelial subtypes. (**N**) Summary schematic illustrating the proposed consequences of pericyte loss on cECs, microglia, and white matter integrity. Pericyte depletion induces endothelial reprogramming characterised by reduced MFSD2A expression, increased VCAM-1 expression, activation of type I interferon signalling (including *Isg15* and *Ifit3*), and shifts in transcription factor activity with reduced SMAD-associated programmes and increased IRF/NF-_κ_B signalling. These changes are associated with microglial activation, myelin basic protein (MBP) loss, and white matter injury relevant to human SVD. For scRNA-seq, n = 4 mice per group (5d CT, 5d DT, 30d CT, and 30d DT). Immunofluorescence statistics were performed using linear mixed-effects models with Bonferroni correction (***, p < 0.001; *, p < 0.05; ^#^, p = 0.05-0.1 - with treatment and timepoint as fixed effects and mouse ID as a random effect).

Differential expression analysis in cECs at 5 days post-depletion revealed upregulation of genes associated with endothelial activation (e.g., *Vcam1*) and interferon signalling (including *Isg15, Ifit3, Oasl2*, and *Ifitm3*), alongside downregulation of BBB-associated genes such as *Mfsd2a* (**Fig. 5C**). Pathway analysis confirmed enrichment of type I interferon signalling, antiviral response pathways, cytokine-mediated signalling, and leukocyte adhesion pathways, consistent with an activated and pro-inflammatory endothelial state (**Fig. 5D**).

Consistent with transcriptomic findings, immunohistochemistry demonstrated reduced MFSD2A signal in capillary endothelium (-19%), indicative of BBB dysfunction, and markedly increased VCAM-1 coverage (+434%), consistent with endothelial activation and a shift towards a venous-like phenotype (**Fig. 5E-H**). In parallel, microglial activation was observed, with no significant change in overall Iba1-positive cell density, but a significant increase in Iba1/MHCII double-positive cells (+32.9%), indicating activation of microglia (**Fig. 5I-K**).

Transcription factor activity analysis revealed a marked shift in endothelial regulatory programmes following pericyte depletion (**Fig. 5L**). Interferon- and inflammation-associated transcription factors, including Irf1, Stat2, Irf2, Irf9, Irf4, Irf3, and Nfkb1, showed increased activity, alongside additional regulators such as Brca1, Spi1, and Hdac7. In contrast, several transcription factors associated with vascular homeostasis and endothelial identity, including Smad1, Smad2, Epas1, Mef2a, Hnf1b, and Egr1, exhibited reduced activity. This pattern is consistent with a transition towards an activated endothelial state.

Finally, GSEA revealed significant enrichment of WMH-associated SVD gene signatures across arterial, capillary, and venous endothelial populations at 30 days post-ablation, whereas enrichment of PVS- or AD-associated gene sets was not significant (**Fig. 5M**). The delayed emergence of WMH-associated endothelial signatures paralleled the delayed white matter abnormalities observed following pericyte depletion, including MBP loss and diffusion MRI changes. Interestingly, AD-associated gene sets displayed an opposite trend in NES at 5 and 30 days compared with SVD-associated signatures, although these changes did not reach statistical significance.

Together, these findings support a model in which pericyte loss induces capillary endothelial reprogramming characterised by BBB dysfunction, inflammatory activation, interferon signalling, and venular-like endothelial transitions that ultimately contribute to white matter injury and human SVD-related molecular states (**Fig. 5N**).

## Discussion

In the present study, we combined an *Atp13a5*-driven genetic strategy, advanced microscopy, ML-based morphometric analyses, longitudinal MRI, and scRNA-seq to investigate the role of brain pericytes in cerebrovascular homeostasis and brain health. Atp13a5 represents one of the most CNS-enriched pericyte markers currently available and enables selective targeting of brain mural cells with no peripheral involvement^16^. Using this mouse model, we demonstrate that moderate pericyte depletion induced transient cerebrovascular dysfunction, endothelial inflammatory remodelling, and delayed white matter alterations associated with transcriptional programmes relevant to human cerebral SVD.

Pericytes and other mural cells are increasingly recognised as highly heterogeneous populations with distinct anatomical locations, morphologies, and functions along the vascular tree^6,13^. However, defining discrete mural cell populations remains challenging, as many canonical markers are shared across multiple vascular cell types. Markers commonly used to identify pericytes, including *Kcnj8, Abcc9, Anpep*, and *Pdgfrb*, are not fully restricted to CNS capillary pericytes and can also label other mural cell populations. Our transcriptomic analyses support the concept that mural cells exist along a transcriptional and anatomical continuum rather than as strictly segregated cell populations. While *Atp13a5* showed strong enrichment in CNS pericytes, we also detected expression in selected vSMC-associated populations at both the transcriptomic and histological levels. Thus, although *Atp13a5* appears substantially more CNS- and pericyte-selective than NG2- or PDGFRβ-based approaches, its expression outside classical capillary “bump-on-a-log” pericytes should still be considered when interpreting depletion studies or genetic manipulations.

To our knowledge, this is the first study to combine high-resolution vascular imaging with ML-based classification to characterise pericyte subtype organisation and remodelling across the brain microvasculature. Using this framework, we identified distinct ensheathing, mesh, and thin-strand pericyte morphotypes distributed along the vascular hierarchy, building on prior morphological classifications of cortical mural cells and pericyte subtypes^15^. Importantly, these subpopulations appeared differentially affected by pericyte depletion, supporting the possibility that specific mural cell populations display distinct vulnerabilities or functions. Although the precise physiological roles of these morphotypes remain incompletely understood, the observed remodelling following ablation suggests substantial plasticity within the pericyte network.

Compared with several previous pericyte-ablation models^9,28^, the extent of pericyte depletion observed in the present study was relatively modest and heterogeneous. This may initially appear as a limitation; however, it may better recapitulate the partial and progressive pericyte dysfunction observed in ageing and human SVD and AD^7,10,11,29^, where complete mural cell ablation is unlikely to occur. Importantly, although pericyte soma density was significantly reduced, overall pericyte coverage was comparatively less affected. This discrepancy strongly suggests that the pericyte network remains highly dynamic, with neighbouring pericytes potentially extending their processes to compensate for local cell loss, as previously proposed by Shih and colleagues^30^. Such compensatory remodelling may contribute to partial preservation of vascular structure despite substantial reductions in pericyte cell numbers.

Interestingly, vascular structural alterations remained relatively limited overall, despite measurable pericyte loss. However, selective increases in vascular diameter were observed in the hippocampus and thalamus during the recovery phase. These regional changes may reflect compensatory vascular adaptation or reduced vascular tone following mural cell dysfunction. Importantly, pericyte numbers returned close to baseline levels by 30 days post-ablation, accompanied by recovery of CBF and BBB permeability. The origin of these repopulating cells remains unclear and warrants further investigation. Potential mechanisms may include proliferation of surviving pericytes, endothelial-to-mesenchymal transition (EndoMT), recruitment of fibroblast-like perivascular cells, or contributions from immune-associated populations. Interestingly, pericyte recovery after depletion has also been reported in cardiac context, supporting the broader concept that mural cell populations can regenerate or repopulate after injury, although the cellular source remains unresolved^31^.

Functionally, the most prominent acute consequence of pericyte depletion was reduced CBF. This finding supports the concept that microvascular pericytes are critical regulators of capillary perfusion and vascular resistance. Notably, CBF deficits occurred despite relatively preserved overall pericyte coverage, consistent with previous work suggesting that contractile regulation is primarily mediated by the pericyte soma rather than distal cellular processes^32,33^. The observed increase in ensheathing pericytes may therefore represent compensatory remodelling aimed at restoring vascular resistance and perfusion, although this interpretation remains speculative.

Blood-brain barrier dysfunction appeared more subtle and delayed than CBF changes, with significant increases in Gadolinium permeability emerging at day 15. This suggests that different aspects of cerebrovascular homeostasis may exhibit distinct sensitivities to pericyte dysfunction. Reduced MFSD2A expression further supports impaired suppression of endothelial transcytosis following pericyte loss. As MFSD2A is a critical regulator of BBB integrity through inhibition of caveolae-mediated transport^34,35^, reduced expression may contribute to increased vesicular trafficking and barrier dysfunction. Different degrees or spatial patterns of pericyte loss may therefore affect BBB integrity through distinct mechanisms, including altered transcytosis, endothelial activation, inflammatory signalling, or junctional disruption.

One particularly interesting finding was the vulnerability of white matter following pericyte depletion. One of the most pronounced reductions in pericyte density was observed in the corpus callosum, a white matter region particularly vulnerable to hypoperfusion and commonly affected in SVD^36^. Diffusion MRI demonstrated subtle but significant white matter alterations, while histological analyses revealed reduced MBP despite preserved overall oligodendrocyte numbers. Together, these findings suggest that moderate pericyte dysfunction may impair white matter integrity without inducing overt oligodendrocyte loss. Increased numbers of OPC-like cells may further indicate ongoing repair or remyelination attempts following vascular injury, consistent with the dynamic interactions between pericytes and OPCs within the perivascular white matter niche^37^. Importantly, white matter abnormalities persisted despite partial recovery of vascular function, suggesting that transient cerebrovascular dysfunction may initiate longer-lasting tissue injury. Reduced CBF, impaired trophic support, BBB dysfunction, and increased exposure to circulating inflammatory factors may together contribute to cumulative white matter damage over time, mechanisms increasingly implicated in vascular cognitive impairment and white matter degeneration^38,39^.

At the molecular level, pericyte loss induced a striking endothelial inflammatory response, particularly within capillary endothelial cells. Among the most prominent changes was the induction of interferon-responsive genes and transcriptional regulators, including *Isg15, Ifit3, Oasl2*, and *Ifitm3*, together with increased activity of IRF-, STAT-, and NFκB-associated transcriptional programmes. These transcriptional regulators are central components of type I interferon and inflammatory signalling cascades and are increasingly recognised as important mediators of endothelial activation and BBB dysfunction in neurological disease^40,41^. Concurrently, endothelial homeostatic regulators and BBB-associated genes such as *Mfsd2a* were reduced. These changes were accompanied by marked upregulation of VCAM-1 and activation of MHCII-positive microglia. Together, these findings suggest that pericytes actively suppress endothelial inflammatory activation under homeostatic conditions.

Although interferon signalling in brain endothelial cells remains incompletely understood in the context of dementia, increasing evidence suggests that type I interferon pathways influence BBB function, vascular inflammation, and immune-cell interactions within the NGVU^41,42^. Interferon-responsive genes such as *Ifitm3* have also been implicated in AD and neuroinflammatory states^43^. Interestingly, Maë and colleagues similarly observed enrichment of interferon-associated pathways in a genetic model of chronic pericyte deficiency, although this pathway was not explored mechanistically^44^. Our findings therefore support the possibility that interferon signalling represents an important downstream consequence of pericyte dysfunction. Whether this inflammatory programme directly contributes to capillary dysfunction, immune-cell adhesion, BBB impairment, or white matter injury remains to be determined.

The observed increase in VCAM-1 expression may be particularly relevant functionally. Under physiological conditions, VCAM-1 expression is predominantly associated with venous endothelial cells and is typically low or absent in capillary endothelial cells^13^. Its robust induction following pericyte loss therefore further supports the concept of a partial venous-shifted endothelial identity within the capillary bed^11^. VCAM-1 expression within capillary endothelial cells could increase leukocyte adhesion and capillary stalling, thereby contributing to impaired microvascular perfusion^45^. The robust induction of VCAM-1 following pericyte loss is also particularly interesting considering recent plasma proteomic work showing that VCAM-1 is among the top endothelial-derived molecules increased during ageing^46^ and associated with imaging markers of cerebral SVD^47^. Concurrently, the reduction in MFSD2A and enrichment of venous-associated transcriptional programmes suggest that capillary endothelial cells undergo partial loss of capillary identity and transition towards a more activated, venular-like endothelial state following pericyte depletion. Similar endothelial phenotypic shifts have been experimentally reported in other contexts of vascular dysfunction, including chronic pericyte deficiency in PDGFRβ mutant mice^44^ and inflammatory BBB disruption in experimental autoimmune encephalomyelitis, where capillary endothelial cells acquire activated and venous-associated transcriptional features^48^.

Finally, comparison of our endothelial transcriptomic data with human GWAS datasets revealed significant enrichment of WMH-associated SVD genes across arterial, capillary, and venous endothelial populations at 30 days post-ablation. In contrast, enrichment of PVS- or AD-associated gene sets was not significant^49^. The delayed emergence of WMH-associated signatures is particularly interesting given the similarly delayed white matter abnormalities observed in our model. Several overlapping genes, including *Col4a1, Col4a2, Prkch, Nid2*, and *Plekhg1*, have previously been implicated in vascular stability, extracellular matrix integrity, and SVD susceptibility^50,51^. In particular, *COL4A1* and *COL4A2* encode key basement membrane components strongly associated with monogenic and sporadic forms of cerebral SVD, while *PRKCH* and *PLEKHG1* have been linked to WMH burden and vascular dysfunction in large GWAS studies^52,53^. Interestingly, this overlap was most pronounced for WMH rather than PVS or AD gene sets, further supporting a closer relationship between pericyte dysfunction, endothelial remodelling, and white matter pathology. Together, these findings support the concept that moderate pericyte dysfunction may induce endothelial transcriptional states relevant to human SVD and progressive white matter pathology.

Several limitations should nevertheless be acknowledged. The present model induces relatively modest and acute pericyte depletion in young adult mice and may not fully recapitulate the chronic progressive vascular dysfunction associated with ageing or dementia. In addition, Atp13a5 expression is not completely restricted to classical capillary “bump-on-a-log” pericytes, and contributions from other mural cell populations cannot be entirely excluded. Finally, although our data strongly implicate interferon-associated endothelial activation following pericyte loss, direct causal relationships between these pathways and downstream vascular or white matter dysfunction remain to be established.

In summary, the present study identifies brain pericytes as central regulators of cerebrovascular homeostasis, endothelial identity, and white matter integrity. Our findings support a model in which even moderate pericyte dysfunction is sufficient to induce endothelial inflammatory remodelling and vascular states associated with human SVD. These results further highlight the importance of the NGVU in age-related cerebrovascular and neurodegenerative disease.

## Acknowledgements

We thank Dr. Zhen Zhao (Temple University) for generously providing the *Atp13a5*-CreERT2 mouse line used in this study. We also thank the BVS staff, technicians, and veterinary team for their invaluable support and expertise, which ensured that all animal procedures were conducted to the highest possible standards of care and animal welfare.

## Contributions

D.S. performed the *in vivo* experiments, MRI acquisitions, immunofluorescence staining, and data analyses, and contributed to figure preparation. A.C. developed and performed the machine learning analyses, developed analytical tools for microscopy image analysis, contributed to MRI acquisitions and immunofluorescence staining, performed data analyses, and generated most of the figures. M.S. performed all bioinformatic analyses for scRNA-seq and conducted statistical analyses. N.F. performed vascular RNA-seq experiments and contributed to immunofluorescence staining and analyses. C.M.Q. contributed to *in vivo* experiments, tissue collection, and immunofluorescence staining and analyses. O.J.U. contributed to immunofluorescence staining and analyses and assisted with figure preparation. S.B. and M.R.B. contributed to immunofluorescence staining and analyses. W.M. and V.W.G. supported animal work and welfare. T.D.F. performed DTI-MRI analyses. R.L. and M.A.J. contributed to MRI acquisition. O.D. provided support for scRNA-seq and statistical analyses. J.C.W. performed spinning-disk confocal microscopy. A.M. conceived and directed the study and coordinated the collaborative work. A.M. wrote the manuscript with substantial contributions from D.S., A.C., and M.S. All authors reviewed and approved the manuscript.

